# PS4: a Next-Generation Dataset for Protein Single Sequence Secondary Structure Prediction

**DOI:** 10.1101/2023.02.28.530456

**Authors:** Omar Peracha

## Abstract

Protein secondary structure prediction is a subproblem of protein folding. A lightweight algorithm capable of accurately predicting secondary structure from only the protein residue sequence could provide a useful input for tertiary structure prediction, alleviating the reliance on MSA typically seen in today’s best-performing models. Unfortunately, existing datasets for secondary structure prediction are small, creating a bottleneck. We present PS4, a dataset of 18,731 non-redundant protein chains and their respective secondary structure labels. Each chain is identified, and the dataset is also non-redundant against other secondary structure datasets commonly seen in the literature. We perform ablation studies by training secondary structure prediction algorithms on the PS4 training set, and obtain state-of-the-art accuracy on the CB513 test set in zero shots.

## 1. Introduction

Recent years have seen great advances in automated protein structure prediction, with open-sourced algorithms, capable in many cases of matching the accuracy of traditional methods for determining the structure of a folded protein such as x-ray crystallography and cryo-EM, made increasingly available. It remains common for the best-performing approaches to rely on MSA data in order to provide strong results (Jumper et al., 2021; Baek et al., 2021; Zheng et al., 2022). One drawback of this is that these algorithms perform poorly on orphan proteins. Another is that quite significant extra resources are required when running these algorithms, particularly disk space for storing the database of potential homologues and computation time to adequately perform a search through several hundred gigabytes of this data. For example, at the time of writing, the lighter-weight version of AlphaFold2 currently made available by DeepMind as an official release^1^ requires 600 GB of disk space, and comes with accuracy tradeoffs; the full version occupies terabytes.

Improved performance on orphan proteins is particularly desirable as it may open up wider avenues for exploration when it comes to the set of possible human-designed proteins which can benefit from reliable algorithmic structure prediction, in turn offering advantages such as faster drug development. Meanwhile, reducing the resource requirements for using protein structure prediction algorithms increases their accessibility, ultimately improving the rate of research advances and downstream industry adoption.

More recently, accurate structure prediction models have been proposed which do not rely on MSA. Wang et al. (2022) propose trRosettaX-Single, a model designed to predict tertiary structure from single-sequence input. They instead leverage a large transformer-based protein ‘‘language model”, so named because it is an algorithm trained to denoise or autoregressively predict residues in a sequence of protein amino acids in a self-supervised manner, much as language models are trained to denoise or autoregressively predict words in a sequence of text (Alaparthi and Mishra, 2020; Radford et al., 2019). This technique has gained popularity because the embeddings generated by the trained model in response to a protein sequence input seem to encode information regarding genetic relationships which a downstream neural network can take advantage of. Furthermore, these models can be trained on the residue data alone, without requiring further labels such as atomic coordinates.

To form one component of trRosettaX-Single, Wang et al. (2022) also use knowledge distillation from a pre-trained MSA-based network (Hinton et al., 2015), training a smaller ‘student” neural network to approximate the MSA-based “teacher” network’s output distribution when the former is fed only a single sequence input, as a way to further induce some understanding of the homology relationships in their model. While performance is close to AlphaFold2 when evaluating structure prediction accuracy on a dataset of human-designed proteins, and exceeds it on a dataset of orphan proteins, the authors point out that accuracy on those orphan proteins is still far from satisfactory. However, the use of upstream neural networks, such as the protein language model, in place of searching large databases, ultimately reduces the resource requirement compared to AlphaFold2.

It is a well-held belief that a protein’s secondary structure has implications that affect the final fold, for example through a correlation with fold rate in certain conditions (Ji and Li, 2010; Huang et al., 2015). We therefore infer that accurate secondary structure prediction models can also serve as powerful upstream components for tertiary structure prediction algorithms. Secondary structure motifs, comprising just a handful of varieties, occur in most proteins with well-defined tertiary structures; indeed, the same classes of secondary structure can occur in proteins that are evolutionarily distant from each other. Furthermore, a significant proportion of all protein structure is in some form of secondary structure (Kabsch and Sander, 1983).

Secondary structure is influenced to a great degree by the local constitution of a residue chain, particularly in the case of helices and coils, rather than by the idiosyncrasies which begin to emerge over the length of an entire protein chain and in turn contribute to the plethora of topologies observed among fully-folded proteins. The implication is that it may be possible to infer the patterns in polypeptide sequences which correspond to the occurrence of the various classes of secondary structure without relying on homology data. However, attempts to prove this empirically have been hampered by a lack of large, high quality datasets for single sequence secondary structure prediction.

Among the most-cited in the literature is the CB513 dataset (Cuff and Barton, 1999), consisting of 513 protein sequences split into training and validation sets. A training set of a few hundred sequences is not sufficient to achieve high test set accuracy on CB513, therefore a typical approach is to use extra training data (Torrisi et al., 2019; Elnaggar et al., 2022). However, the specific proteins included in CB513 are not identified, which can make it difficult to ensure there are no duplicate occurrences of samples from the test set in the augmented training set. Furthermore, the CB513 test set and training set contain some instances of different subsequences extracted from the same protein chain. Although local information is likely to play a strong role in determining the materialisation of secondary structure motifs, it cannot be said for certain that there is no information leakage between the training and the test set, suggesting evaluation on the CB513 test set is unideal in cases where its training set was also seen by the model. Unfortunately, the lack of large datasets for protein secondary structure prediction hitherto means that omission of the CB513 from a larger training superset would be a significant sacrifice.

The majority of other datasets seen in the literature are of similar or smaller size to CB513 (Drozdetskiy et al., 2015; Yang et al., 2018). Klausen et al. (2019) introduce a notably larger dataset, comprising almost 11,000 highly non-redundant protein sequences. Elnaggar et al. (2022) were able to leverage this dataset, among other smaller ones including the CB513 training set, to achieve a test set Q8 accuracy of 74.5% and Q3 accuracy of 86.0% on the CB513. Not only is this the highest previously reported, albeit by a narrow margin, but the authors were able to avoid using extra inputs such as MSA by leveraging a protein language model. This is the first time accuracy in that range has been achieved without the use of MSA, to our knowledge.

In order to continue to push the boundaries further, we propose PS4, a dataset for protein secondary structure prediction comprising 18,731 sequences. Each sequence is from a distinct protein chain and consists of the entire resolved chain at the time of compilation, including chains featuring multiple domains. Samples are filtered at 40% similarity via a levenshtein distance comparison to ensure a highly diverse dataset. Crucially, samples are also filtered by similarity to the entire CB513 dataset in the same manner, allowing for improved evaluation reliability when using the CB513 for performance benchmarking. All samples are identified by their PDB code and chain ID, and are also guaranteed to have a corresponding entry in the CATH database (Knudsen and Wiuf, 2010), to facilitate research on further hierarchal modelling tasks such as domain location prediction or domain classification.

We perform ablation studies by using the PS4 training set and evaluating on both the PS4 test set and the CB513 test set in a zero-shot manner, leaving out the CB513 training set from the process. We use the same protein language model as Elnaggar et al. (2022) to extract input embeddings and evaluate on multiple neural network architectures for the end classifier, with no further inputs such as MSA. We obtain state-of-the-art results for Q3 and Q8 secondary structure prediction accuracy on the CB513, 86.8% and 76.3% respectively, by training solely on PS4.

We make the full dataset freely available along with code for evaluating our pretrained models, for training them from scratch to reproduce our results and for running predictions on new protein sequences. Finally, in the interests of obtaining a dataset of sufficient scale to truly maximise the potential of representation learning for secondary structure prediction, we provide a toolkit for any researchers to add new sequences to the dataset, ensuring the same criteria for non-redundancy. New additions will be released in labelled versions to ensure the possibility for consistent benchmarking in a future-proof manner^2^.

**Fig. 1:**
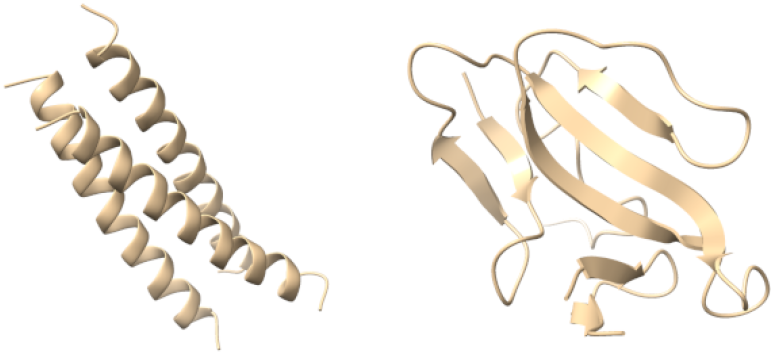
Two folded proteins displaying different common secondary structure motifs, rendered using UCSF ChimeraX software for molecular visualisation (Pettersen et al., 2021). (Left) A synthetic triple-stranded protein by Lovejoy et al. (1993), featuring alpha helices. (Right) A two-chain synthetic structure by Scherf et al. (2001) predominantly featuring beta strands.

## 2. Methods

### 2.1. Dataset Preparation

The PS4 dataset consists of 18,731 protein sequences, split into 17,799 training samples and 932 validation samples, where each sequence is from a distinct protein chain. We first obtained the entire precomputed DSSP database (Kabsch and Sander, 1983; Joosten et al., 2011), initiating the database download on 16th December 2021^3^. The full database at that time contained secondary structure information for 169,579 proteins, many of which are multimers, in DSSP format, with each identified by its respective PDB code.

We iterate through the precomputed DSSP files and create a separate record for each individual chain, noting its chain ID, its residue sequence as a string of one-letter amino acid codes, and its respective secondary structure sequence, assigning one of nine possible secondary structure classes for each residue in the given chain; the ninth class, the polyproline helix, has not generally been taken into consideration by other secondary structure prediction algorithms, and we too ignore this class when performing our own algorithmic assessments, however the information is retained in the raw dataset provided.

We also store the residue number of the first residue denoted in the DSSP file for the chain, which is quite often not number 1; being able to infer the residue number of any given residue in the chain could better facilitate the ability to use external data by future researchers. Following from whichever is the first residue included for that chain, we omit chains which are missing any subsequent residues. We further omit any chains containing fewer than 16 residues. Finally we perform filtration to greatly reduce redundancy, checking for similarity below 40% against the entire CB513 dataset, and then for all remaining samples against each other.

We chose the levenshtein distance to compute similarity due to its balance of effectiveness as a distance metric for biological sequences (Berger et al., 2021), its relative speed and its portability, with optimised implementations existing in several programming languages. This last property is of particular importance when factoring in our aim for the PS4 dataset to be extensible by the community, enabling a federated approach to maximally scale and leverage the capabilities of deep learning.

The possibility of running similarity checks locally with a non-specialised computing setup means that even a bioinformatics hobbyist can add new sequences to future versions of the dataset and guarantee a consistent level of non-redundancy, without relying on precomputed similarity clusters. This removes hurdles towards future growth and utility of the PS4 dataset, while also allowing lightweight similarity measurement against proteins which are not easily identifiable by a PDB or UniProt code, such as those in the CB513.

As a last step, we omit any chains which do not have entries in the CATH database as of 16th December 2021, ruling out roughly 1,500 samples from the final dataset. We make the CATH data of all samples in the PS4 available alongside the dataset, in case future research is able to leverage that structural data for improved performance on secondary structure prediction or related tasks. We ultimately chose to focus the scope of this work purely on predictions from single sequence, and did not find great merit in early attempts to include domain classification within the secondary sequence prediction pipeline; as such, this restriction will not be enforced for community additions to PS4.

The main secondary structure data is made available as a CSV file, which is 8.2 MB in size. The supplemental CATH data is a 1.3 MB file in pickle format, mapping chain ID to a list of domain boundary residue indices and the corresponding four-integer CATH classification. Finally, a file in compressed NumPy format (Harris et al., 2020) maps chain IDs to the training or validation set, according the the split used in our experiments.

### 2.2. Experimental Evaluation

We conduct experiments to validate the PS4 dataset’s suitability for use in secondary structure prediction tasks. We train two models, each based on a different neural network algorithm, to predict eight-class secondary structure given single sequence protein input. The models are trained only on the PS4 training set and then evaluated on both the PS4 test set and the CB513 test set. We avoid using any training data from the CB513 training set, meaning evaluation on its test set is conducted in zero shots. We also do not provide surrounding ground truth data at evaluation time for those samples in the CB513 test set which are masked subsequences of a training sample, but rather predict the secondary structure for the whole sequence at once.

Both models make use of the pretrained, open-source encoder-only version of ProtT5-XL-UniRef50 model by Elnaggar et al. (2022) to generate input embeddings from the initial sequence. Our algorithms are composable such that any protein language model could be used in place of ProtT5-XL-UniRef50, opening up an avenue for potential future improvement, however our choice was in part governed by a desire to maximise accessibility; we use the half-precision version of the model made publicly available by Elnaggar et al. (2022), which can fit in just 8 GB of GPU RAM. As such, our entire training and inferencing pipeline can fit on a single GPU.

The protein language model generates an *N* × 1024 encoding matrix, where *N* is the number of residues in the given protein chain. Our two models differ in the neural network architecture used to form the classifier component of our overall algorithm, which generates secondary structure predictions from these encoding matrices. Our first model, which we call PS4-Mega, leverages 11 moving average equipped gated attention (Mega) encoder layers (Ma et al., 2022) to compute a final encoding which is then passed to an output affine layer and softmax to generate a probability distribution over the eight secondary structure classes for each residue in the sequence.

**Fig. 2:**
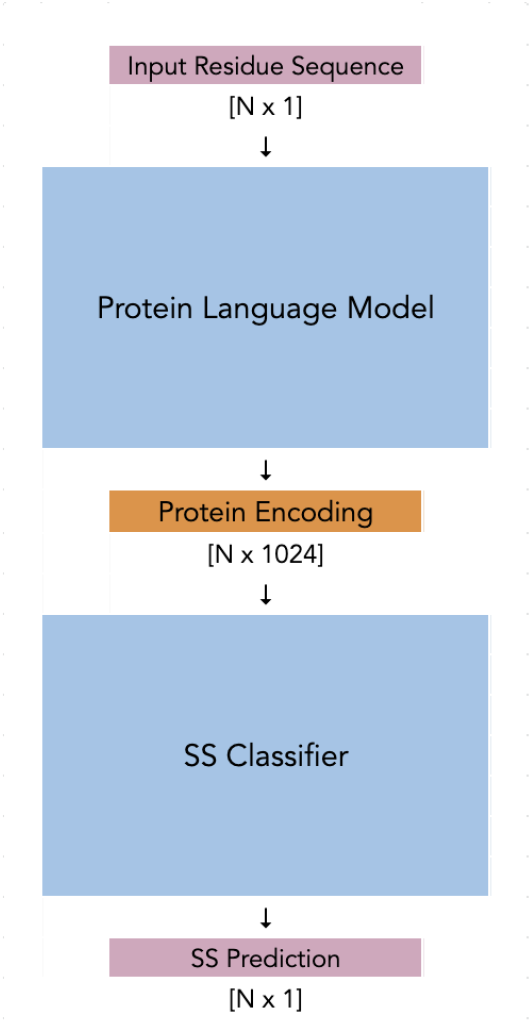
Overview of the neural network meta architecture used in our experiments. The protein language model here refers to an encoder-only half-precision version of ProtT5-XL-UniRef50, while the SS classifier is either a mega-based or convolution-based network. We precompute the protein encodings generated by the pretrained language model, reducing the computations necessary during training to only the forwards and backwards passes of the classifier.

We chose Mega encoders due to their improved inductive bias when compared to a basic transformer (Vaswani et al., 2017), which promises to offer a better balance of factoring both local and global dependencies when encoding the protein sequence. We use a hidden dimension of 1024 for our Mega encoder layers, a z_dim of 128 and an n_dim of 16. We use a dropout of probability 0.1 on the moving average gated attention layers and the normalised feedforward layers, and the simple variant of the relative positional bias. Normalisation is computed via layernorm.

Our second algorithm, which we call PS4-Conv, is derived from the secondary structure prediction model used by Elnaggar et al. (2022) and is entirely based on 2-dimensional convolutional layers. We found that the exact model they used did not have sufficient capacity to fully fit our training set, likely due to it comprising many more samples, and so our convolutional neural classifier is larger, using 5 layers of gradually reducing size, rather than 2. All layers use feature row-wise padding of 3 elements, a 7 × 1 kernel size and a stride of 1. All layers but the last are followed by a ReLU activation and a dropout layer, with probability 0.1. Both models are trained to minimise a multiclass cross entropy objective.

## 3. Implementation

### 3.1. Model Training

All neural network training, evaluation and inference logic is implemented using PyTorch (Paszke et al., 2019). We train both models for 30 epochs, using Adam optimiser (Kingma and Ba, 2014) with 3 epochs of warmup and a batch size of 1, equating to 53,397 warmup steps. Both models increase the learning rate from 10^-7^ to a maximum value of 0.0001, chosen by conducting a learning rate range test (Smith and Topin, 2017), during the warmup phase before gradually reducing back to 10^-7^ via cosine annealing.

The input embeddings from ProtT5-XL-UniRef50 are precomputed, requiring roughly one hour to generate these for the entire PS4 dataset on GPU. Hence only the weights of the classifier component are updated by gradient descent, while the encoder protein language model maintains its original weights from pretraining. For convenience and extensibility, we make a script available in our repository to generate these embeddings from any FASTA file, allowing for predictions on novel proteins.

PS4-Mega has 83.8 million parameters and takes roughly 40 minutes per epoch when training on a single GPU. PS4-Conv is much more lightweight, with just 4.8 million parameters and requiring only 13.5 minutes per epoch on GPU. Both models are trained in full-precision. PS4-Mega obtains a training and test set accuracy of 99.4% and 78.2% on the PS4 dataset for Q8 secondary structure prediction respectively, and 76.3% on CB513. PS4-Conv performs almost as well, obtaining 93.1% training and 77.9% test set accuracy on PS4, and 75.6% on CB513. Furthermore, both algorithms show an improvement over state-of-the-art for Q3 accuracy, as shown in Table 2.

**Table 1.** A comparison of commonly-cited datasets for secondary structure prediction in recent literature. Ours is by far the largest and the only one in which the proteins can be identified. Only ours and CB513 are fully self-contained for training and evaluation with specified training and test sets. The CB513 achieves this distinction in many cases by masking subsequences of training samples, only sometimes including an entire sequence as a whole in the test set.

| Dataset | Samples | Train/Test Split | Has PDB Codes |
| --- | --- | --- | --- |
| TS115 | 115 | N/A | No |
| NEW364 | 364 | N/A | No |
| CB513 | 511 | via masking | No |
| NetSurfP-2.0 | 10792 | N/A | No |
| **PS4** | **18731** | **17799 / 932** | **Yes** |

**Table 2.**
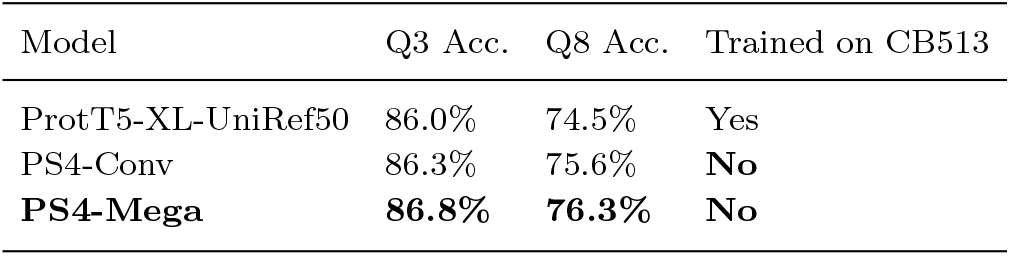
A comparison of Q3 and Q8 performance on the CB513 test set by the leading algorithms for secondary structure prediction, all of which use the same protein language model and operate without MSA input. The version of ProtT5-XL-UniRef50 shown here includes a convolution-based classifier network. Results for ProtT5-XL-UniRef50 are quoted directly from Elnaggar et al. (2022).

### 3.2. Dataset Extension

Our initial dataset preprocessing, from which the secondary structure CSV was obtained, was implemented in the Rust programming language. Filtering through over 160,000 proteins in Python was prohibitively slow. In particular, performing that many string comparisons to verify non-redundancy runs in O(*n*^2^) time complexity, and given a large value of *n* as seen in our case, the speed advantages offered by Rust and its various features were necessary to be able to complete preprocessing on a simple computing setup, running only on a quad-core Intel i5 CPU. We were able to leverage multithreading when iterating through over 169k DSSP files, and could therefore increase the efficiency of the sequence comparisons via a parallelised divide-and-conquer approach, reducing time complexity to O(*n* × *log*(*n*)).

Since all training, evaluation and model inference code is made available in a Python library, it would be convenient for all data processing code to also be Python-based. Therefore, we make the original Rust code callable via Python, such that anyone wishing to add new samples to a future version of PS4 can still leverage the original implementation for similarity measurement, ensuring the quality of the dataset is sustainable. Even with the optimisations made, preprocessing the original dataset on common commercial hardware still required close to 2 days. Fortunately, smaller future additions which comprise far fewer than 169k sequences will run faster, from seconds to minutes, due to the rapidly-growing nature of superlinear time complexity.

Initial extensions will be made via pull request to the maintained code repository for the dataset. Sufficient added sequences will give rise to a new, versioned release of the dataset. Future improvements to the PS4 may seek to further simplify the process for community-led extension, for example managed via a web graphic user interface, so as to maximise accessibility.

## 4. Discussion

We have presented the largest dataset for secondary structure prediction and made available a pipeline for further growth of the dataset by the bioinformatics community. The promise of learning-based algorithms to be a catalyst of progress on tasks related to protein folding is significant, particularly given what has been seen in recent advances in tertiary structure structure prediction. However, for this promise to be realised requires datasets of sufficient scale and quality.

The state of datasets for protein secondary structure prediction has been such that most recent advances in the literature have depended on an amalgamation of different sources of data for both the training and evaluation sets in order to maximise the number of samples. This instantly hampers progress by creating an obstacle towards the acquisition of good quality data by would be researchers, as well as well-attested benchmarks to measure against. Because proteins sequences in pre-existing datasets have typically been difficult to identify, reliability of assessments may also be an issue, resulting from the possibility of leakage between training and evaluation data.

The most common method to mitigate this issue so far has been using a cutoff date threshold, for example only evaluating an algorithm on proteins released after a date known to be after all samples in the training set were themselves released. This has the downside of either limiting future research to the same datasets, which are still too small to fully maximise the potential offered by deep learning algorithms, or in the case that new datasets are introduced in future, immediately invalidating the evaluation data used in setting a previous benchmark.

We show that by training on the PS4 dataset, we can achieve new state-of-the-art performance on the CB513 test set, validated on multiple classifier architectures. Our method is composable such that alternative protein language models can be used to generate embeddings, should this prove useful to future researchers. We also impose strict sequence similarity restrictions and run these directly against the CB513 dataset as a whole to greatly reduce the probability of data leakage into the test set, with respect to both the CB513 and the PS4’s own validation set.

At the same time, we acknowledge that the PS4 is still too small to truly support the development of a general learning-based solution to protein single sequence secondary structure prediction.

Therefore, we chose to leverage the scaling opportunities offered by open-source technology and provided a protocol for the community to continue augmenting the dataset with new samples. Given a file in PDB format, with full atomic co-ordinates, it is trivial to assign secondary structure to each residue using DSSP. As the PDB continues to grow, and indeed with the arrival of new protein structure databases with atomic co-ordinates resolved via automated methods of increasingly high quality (Varadi et al., 2021), the task of single sequence secondary structure prediction should be able to benefit from increased data availability over time and positively feed back into the cycle by supporting the improvement of tertiary structure prediction algorithms in turn.

We propose the PS4 dataset as a hub for protein secondary structure data for training learning algorithms; a common first point of call where researchers can reliably obtain a high quality dataset and benchmark against other algorithms with confidence of that data’s cleanliness. To achieve this, making it able to grow as new labelled data becomes available is a first step. Future developments could focus on user experience and quality of life improvements to better facilitate community contributions, thus maximising overall effectiveness.

## Footnotes

1 Available at https://github.com/deepmind/alphafold

2 Available at https://github.com/omarperacha/ps4-dataset

3 Available by following the instructions at https://swift.cmbi.umcn.nl/gv/dssp/DSSP1.html

## References

S. Alaparthi and M. Mishra. Bidirectional encoder representations from transformers (bert): A sentiment analysis odyssey, 2020. URL https://arxiv.org/abs/2007.01127.

M. Baek, F. DiMaio, I. Anishchenko, J. Dauparas, S. Ovchinnikov, G. R. Lee, J. Wang, Q. Cong, L. N. Kinch, R. D. Schaeffer, C. Millan, H. Park, C. Adams, C. R. Glassman, A. DeGiovanni, J. H. Pereira, A. V. Rodrigues, A. A. van Dijk, A. C. Ebrecht, D. J. Opperman, T. Sagmeister, C. Buhlheller, T. Pavkov-Keller, M. K. Rathinaswamy, U. Dalwadi, C. K. Yip, J. E. Burke, K. C. Garcia, N. V. Grishin, P. D. Adams, R. J. Read, and D. Baker. Accurate prediction of protein structures and interactions using a three-track neural network. Science, 373(6557):871–876, 2021.

B. Berger, M. S. Waterman, and Y. W. Yu. Levenshtein distance, sequence comparison and biological database search. IEEE Trans Inf Theory, 67(6):3287–3294, Jun 2021. ISSN 0018-9448 (Print); 0018-9448 (Linking).

J. A. Cuff and G. J. Barton. Evaluation and improvement of multiple sequence methods for protein secondary structure prediction. Proteins, 34(4):508–519, Mar 1999. ISSN 0887-3585 (Print); 0887-3585 (Linking).

A. Drozdetskiy, C. Cole, J. Procter, and G. J. Barton. JPred4: a protein secondary structure prediction server. Nucleic Acids Research, 43(W1):W389–W394, 04 2015. ISSN 0305-1048. doi: 10.1093/nar/gkv332. URL https://doi.org/10.1093/nar/gkv332.

A. Elnaggar, M. Heinzinger, C. Dallago, G. Rehawi, Y. Wang, L. Jones, T. Gibbs, T. Feher, C. Angerer, M. Steinegger, D. Bhowmik, and B. Rost. Prottrans: Toward understanding the language of life through self-supervised learning. IEEE Transactions on Pattern Analysis and Machine Intelligence, 44(10):7112–7127, 2022.

C. R. Harris, K. J. Millman, S. J. van der Walt, R. Gommers, P. Virtanen, D. Cournapeau, E. Wieser, J. Taylor, S. Berg, N. J. Smith, R. Kern, M. Picus, S. Hoyer, M. H. van Kerkwijk, M. Brett, A. Haldane, J. F. del Rio, M. Wiebe, P. Peterson, P. Gérard-Marchant, K. Sheppard, T. Reddy, W. Weckesser, H. Abbasi, C. Gohlke, and T. E. Oliphant. Array programming with NumPy. Nature, 585(7825):357–362, Sept. 2020. doi: 10.1038/s41586-020-2649-2. URL https://doi.org/10.1038/s41586-020-2649-2.

G. Hinton, O. Vinyals, and J. Dean. Distilling the knowledge in a neural network, 2015. URL https://arxiv.org/abs/1503.02531.

J. T. Huang, T. Wang, S. R. Huang, and X. Li. Prediction of protein folding rates from simplified secondary structure alphabet. J Theor Biol, 383:1–6, Oct 2015. ISSN 1095-8541 (Electronic); 0022-5193 (Linking). doi: 10.1016/j.jtbi.2015.07.024.

Y.-Y. Ji and Y.-Q. Li. The role of secondary structure in protein structure selection. The European Physical Journal E, 32(1): 103–107, 2010. doi: 10.1140/epje/i2010-10591-5. URL https://doi.org/10.1140/epje/i2010-10591-5.

R. P. Joosten, T. A. H. te Beek, E. Krieger, M. L. Hekkelman, R. W. W. Hooft, R. Schneider, C. Sander, and G. Vriend. A series of pdb related databases for everyday needs. Nucleic Acids Res, 39(Database issue):D411–9, Jan 2011. ISSN 1362-4962 (Electronic); 0305-1048 (Print); 0305-1048 (Linking). doi: 10.1093/nar/gkq1105.

J. Jumper, R. Evans, A. Pritzel, T. Green, M. Figurnov, O. Ronneberger, K. Tunyasuvunakool, R. Bates, A. žídek, A. Potapenko, A. Bridgland, C. Meyer, S. A. A. Kohl, A. J. Ballard, A. Cowie, B. Romera-Paredes, S. Nikolov, R. Jain, J. Adler, T. Back, S. Petersen, D. Reiman, E. Clancy, M. Zielinski, M. Steinegger, M. Pacholska, T. Berghammer, S. Bodenstein, D. Silver, O. Vinyals, A. W. Senior, K. Kavukcuoglu, P. Kohli, and D. Hassabis. Highly accurate protein structure prediction with alphafold. Nature, 596(7873):583–589, 2021.

W. Kabsch and C. Sander. Dictionary of protein secondary structure: pattern recognition of hydrogen-bonded and geometrical features. Biopolymers, 22(12):2577–2637, Dec 1983.

D. P. Kingma and J. Ba. Adam: A method for stochastic optimization. CoRR, abs/1412.6980, 2014.

M. S. Klausen, M. C. Jespersen, H. Nielsen, K. K. Jensen, V. I. Jurtz, C. K. Sønderby, M. O. A. Sommer, O. Winther, M. Nielsen, B. Petersen, and P. Marcatili. Netsurfp-2.0: Improved prediction of protein structural features by integrated deep learning. Proteins, 87(6):520–527, Jun 2019. ISSN 1097-0134 (Electronic); 0887-3585 (Linking).

M. Knudsen and C. Wiuf. The cath database. Hum Genomics, 4(3):207–212, Feb 2010. ISSN 1479-7364 (Electronic); 1473-9542 (Print); 1473-9542 (Linking). doi: 10.1186/1479-7364-4-3-207.

B. Lovejoy, S. Choe, D. Cascio, D. K. McRorie, W. F. DeGrado, and D. Eisenberg. Crystal structure of a synthetic triple-stranded alpha-helical bundle. Science, 259(5099):1288–1293, Feb 1993. ISSN 0036-8075 (Print); 0036-8075 (Linking). doi: 10.1126/science.8446897.

X. Ma, C. Zhou, X. Kong, J. He, L. Gui, G. Neubig, J. May, and Z. Luke. Mega: Moving average equipped gated attention. arXiv preprint arXiv:2209.10655, 2022.

A. Paszke, S. Gross, F. Massa, A. Lerer, J. Bradbury, G. Chanan, T. Killeen, Z. Lin, N. Gimelshein, L. Antiga, A. Desmaison, A. Kopf, E. Yang, Z. DeVito, M. Raison, A. Tejani, S. Chilamkurthy, B. Steiner, L. Fang, J. Bai, and S. Chintala. Pytorch: An imperative style, high-performance deep learning library. In Advances in Neural Information Processing Systems 32, pages 8024–8035. Curran Associates, Inc., 2019.

E. F. Pettersen, T. D. Goddard, C. C. Huang, E. C. Meng, G. S. Couch, T. I. Croll, J. H. Morris, and T. E. Ferrin. Ucsf chimerax: Structure visualization for researchers, educators, and developers. Protein Sci, 30(1):70–82, Jan 2021. ISSN 1469-896X (Electronic); 0961-8368 (Print); 0961-8368 (Linking). doi: 10.1002/pro.3943.

A. Radford, J. Wu, R. Child, D. Luan, D. Amodei, and I. Sutskever. Language models are unsupervised multitask learners. 2019.

T. Scherf, R. Kasher, M. Balass, M. Fridkin, S. Fuchs, and E. Katchalski-Katzir. A beta-hairpin structure in a 13-mer peptide that binds alpha -bungarotoxin with high affinity and neutralizes its toxicity. Proc Natl Acad Sci U S A, 98(12):6629–6634, Jun 2001. ISSN 0027-8424 (Print); 1091-6490 (Electronic); 0027-8424 (Linking). doi: 10.1073/pnas.111164298.

L. N. Smith and N. Topin. Super-convergence: Very fast training of neural networks using large learning rates, 2017. URL https://arxiv.org/abs/1708.07120.

M. Torrisi, M. Kaleel, and G. Pollastri. Deeper profiles and cascaded recurrent and convolutional neural networks for state-of-the-art protein secondary structure prediction. Scientific Reports, 9(1):12374, 2019. URL https://doi.org/10.1038/s41598-019-48786-x.

M. Varadi, S. Anyango, M. Deshpande, S. Nair, C. Natassia, G. Yordanova, D. Yuan, O. Stroe, G. Wood, A. Laydon, A. žídek, T. Green, K. Tunyasuvunakool, S. Petersen, J. Jumper, E. Clancy, R. Green, A. Vora, M. Lutfi, M. Figurnov, A. Cowie, N. Hobbs, P. Kohli, G. Kleywegt, E. Birney, D. Hassabis, and S. Velankar. AlphaFold Protein Structure Database: massively expanding the structural coverage of protein-sequence space with high-accuracy models. Nucleic Acids Research, 50(D1):D439–D444, 11 2021. ISSN 0305-1048. doi: 10.1093/nar/gkab1061. URL https://doi.org/10.1093/nar/gkab1061.

A. Vaswani, N. Shazeer, N. Parmar, J. Uszkoreit, L. Jones, A. N. Gomez, L. Kaiser, and I. Polosukhin. Attention is all you need. In Proceedings of the 31st International Conference on Neural Information Processing Systems, NIPS’17, pages 6000–6010, Red Hook, NY, USA, 2017. Curran Associates Inc. ISBN 9781510860964.

W. Wang, Z. Peng, and J. Yang. Single-sequence protein structure prediction using supervised transformer protein language models. Nature Computational Science, 2(12):804–814, 2022.

Y. Yang, J. Gao, J. Wang, R. Heffernan, J. Hanson, K. Paliwal, and Y. Zhou. Sixty-five years of the long march in protein secondary structure prediction: the final stretch? Briefings in Bioinformatics, 19(3):482–494, May 2018. ISSN 1477-4054 (Electronic); 1467-5463 (Print); 1467-5463 (Linking). doi: 10.1093/bib/bbw129.

W. Zheng, Q. Wuyun, and P. L. Freddolino. D-i-tasser: Integrating deep learning with multi-msas and threading alignments for protein structure prediction. 15th Community Wide Experiment on the Critical Assessment of Techniques for Protein Structure Prediction, December 2022.

